# CAR T cell cytotoxic responses are rapidly generated and sensitive to the unligated TCR

**DOI:** 10.1101/2025.07.25.666844

**Authors:** Kevin L. Scrudders, Suriya Selvarajan, Kenneth Rodriguez-Lopez, Weichuan Luo, Bo Huang, Suilan Zheng, Geoffrey H. Graff, Francisco N. Barrera, Philip S. Low, Shalini T. Low-Nam

## Abstract

Chimeric antigen receptor (CAR) T cells expressing tumor-targeting engineered receptors can robustly eliminate cancer cells through secretion of cytotoxic factors. Durable remission in leukemia and lymphoma treatment has not been matched in solid tumors. Efforts to maximize tumor destruction and minimize toxicities have driven efforts to tune CAR signaling. However, the molecular mechanisms for CAR triggering and thresholds for activation are incompletely understood. Here, we measured the collection of CAR binding interactions that culminate in polarized delivery of lytic granules to the junction with the target. CAR T cells binarized cytotoxic activities in response to a few binding events and population outcomes were dominated by a subset of cells. Activation at the single molecule level matches the sensitivity of the native T cell receptor (TCR) and points to potent downstream signal propagation. Disruption of the unligated TCR with a transmembrane-targeting inhibitory peptide strongly dampened CAR T cell activation, indicating a critical crosstalk between the two receptors. Harnessing CAR T cell efficacy and reduction of toxicity will require new approaches to modify integration of the binding events, collected stochastically, that are rapidly digitized. These sensitive CAR T cell responses provide new insights into driving cytotoxic signaling through surface interaction engineering.

---

Adoptive transfer of chimeric antigen receptor (CAR) T cells has successfully mobilized anti-tumor cytotoxicity toward cell destruction and demonstrated efficacy in hematologic cancers^1^. Reprogramming cytotoxic T cells to deliver destructive payloads, via granule polarization, to tumor targets can generate potent damage^2–4^. Tumor-associated antigens (TAAs) are frequently overexpressed on tumor targets but are rarely exclusive and expression profiles change during disease progression^5^. CAR T cells have been inefficacious in the treatment of solid tumors, prompting extensive investigation into their mechanisms of activation and opportunities for alternate activation modes or orthogonal control over CAR activation^6–8^. Poor performance has been ascribed to weak signal propagation upon receptor triggering^9–11^ and alterations in the prototypical organization of the intermembrane junction with the target cell^12^. Recent longitudinal data in leukemia patients with sustained responses of more than a decade showed that a subset of clonal CAR T cells dominated the response and, thus, point to the potency of particular cells within the population^1^.

Native T cells are the standout example of cells that collect inputs stochastically. During immune surveillance, single T cells are tuned to respond to very low, single molecule levels of inputs since agonist ligands, typically of low affinities, may be present at low densities on antigen presenting cells^13–19^. In the native context, small numbers of agonist peptide-loaded major histocompatibility complexes (pMHC) bound to T cell receptors (TCRs) are sufficient to mobilize global responses in cytotoxic T cells (CTLs) to secrete cytokines and cytolytic factors, respectively^14,15,20^. Lethal hits to targets via polarization and exocytosis of vesicles laden with perforins and granzymes occur within a few minutes after initial contact^21,22^. Robust signaling appears to be lost in the CAR T cell context, although rapid vesicle polarization has been measured^23^.

In this study, we measured the collection of binding interactions by CAR T cells that map to polarization and degranulation outcomes. A single molecule impulse-response (I-R) assay directly visualizes the stochastic sequence of binding events between ligands and receptors and maps these discrete inputs to global, binarized outcomes. We focus on a CAR T cell that expresses an anti-FITC CAR in these studies. To promote engagement of this anti-FITC CAR T cell with a cancer cell, we use a folate-fluorescein bispecific adapter, termed EC-17, to form a bridge between the anti-FITC CAR on the T cell and a folate receptor on a cancer cell. EC-17 was considered a validated bispecific adapter for this purpose, since it has been repeatedly shown to image multiple human tumors during cancer surgeries^24^ and because it has also been demonstrated to mediate eradication of solid tumors in numerous animal models^25^. Complexes of FRα:EC-17 bridge:anti-FITC CAR were mapped to CAR T cell polarization and showed the prototypical, ensemble, low sensitivity with a threshold agonist density of hundreds of molecules per square micron. However, polarization and cytotoxic degranulation were readily observed at lower ligand densities. Rare CAR T cells that activated at less than 1 molecule per square micron agonist densities maintained the rapid activation phenotype despite experiencing spatially isolated engagement events. The analog-to-digital conversion was independent of ligand density and points to a fundamental retention of single molecule sensitivity in the CAR T cell signaling response. The uncanny resemblance to the native TCR prompted us to explore a direct crosstalk between the receptors. A recently-developed peptide-based inhibitor of the TCR largely abrogated CAR T cell activation, in the absence of TCR ligands. Overall, these results reveal that, despite fundamental differences, CAR T cells engage the intracellular machinery very efficiently.

## Single CAR T cell potency

Early CAR T cell activation culminates in release of cytotoxic factors into the intermembrane junction with the target. We used a second-generation, universal CAR T cell that recognizes the ligand FITC and retains expression of the native T cell receptor (TCR) (Fig. 1a). Expression of the receptor was coupled to GFP expression via a self-cleaving peptide system^26^. The tumor-associated antigen (TAA) folate receptor (FRα) is recognized using an adaptor molecule of FITC linked to folic acid, called EC-17 (Fig. 1a). We characterized anti-FITC CAR T cell killing of MDA-MB-231 cell targets using an *in vitro* cell:cell killing assay. CAR T cells were exposed to targets at a ratio of 4:1 and with saturating EC-17 in solution. Interactions between CAR T cells and MD-MB-231 cells were imaged for up to 16 hours in a diascopic configuration and approximately 50% killing was achieved (Fig. 1b; Movie S1). The tumor cells expressed FRα over a range spanning 2 orders of magnitude and CAR expression on the T cells covered a similar range (Fig. 1c-d). The number of CAR T cell contacts that led to target killing was bimodal with one third of targets being eliminated with fewer than 3 CAR T cell contacts. (Fig. 1e). Those target cells that were killed as the result of larger numbers of contacts may have experienced less productive engagement events or the tumor cells had inherent resilience to destruction^27^. Nonetheless, the ability of some CAR T cells to cause potent damage at the single cell level prompted us to directly measure the activation process for individual cells.

**Figure 1.**
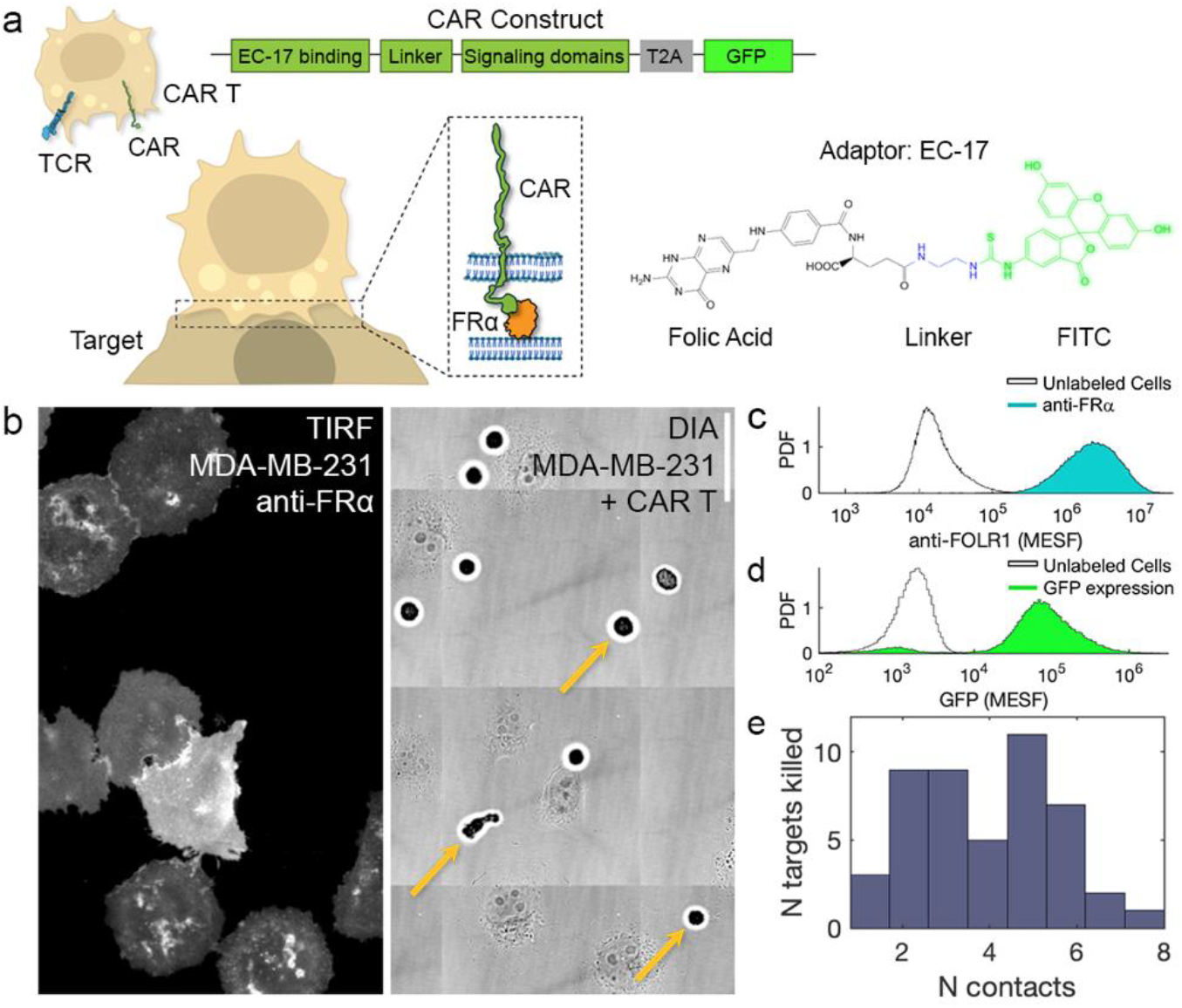
Tumor targets are potently eliminated by small numbers of CAR T cell contacts. **a.** Schematic of CAR T cells expressing both the native TCR and a second-generation anti-FITC CAR. The CAR construct included a self-cleaving T2A sequence and a GFP reporter as a measure of CAR expression. The CAR T cells engage tumor targets at an intermembrane junction. Recognition of tumor targets expressing the antigen FRα is mediated by the bispecific adaptor, EC-17, a bridge molecule comprised of folic acid ligand, a linker sequence, and FITC. **b**. In cell-cell killing experiments, MDA-MB-231 cells were plated on a glass coverslip. Staining with a fluorescently-conjugated anti-FRα, viewed in TIRF, showed a range of expression levels (left). Targets were exposed to CAR T cells at a 4:1 effector:target ratio (right) and monitored using diascopic imaging for up to 8 hours. Yellow arrows indicate example CAR T cells. Scale bar = 20 microns. **c**. Expression of FRα, labeled with anti-FRα and measured by flow cytometry. **d**. Expression of CAR on T cells based on labeling with EC-17 conjugated to an Alexa647 and measured by flow cytometry. **e**. Numbers of CAR T cell contacts that resulted in the death of a target cell.

Reconstitution of immune cell interfaces has enabled unprecedented detail of the binding kinetics and organization that govern these junctions^16,28–39^. When applied to CAR T cells, reconstituted surfaces have provided evidence for weak signaling from ligated CARs, altered organization within the interface, and high activation thresholds^9,23^. We prepared SLBs to mimic the tumor surface with FRα TAA densities ranging from physiological to malignant values (<1/μm^2^ to 100s/μm^2^; Fig. 2a-d; Fig. S1^26^). Mobile FRα, loaded with EC-17 were displayed on supported lipid bilayer (SLB) surfaces in the presence of a high density of the integrin adhesion ligand ICAM, which facilitates CAR T cell landing and spreading. FRα molecules were singly labeled with an organic fluorophore to enable TIRF-based detection and quantification of the CAR ligands on the surface (Fig. 2b-c; Fig. S1) and tracking of receptor:ligand complexes upon engagement of CAR T cells with the surface.

**Figure 2.**
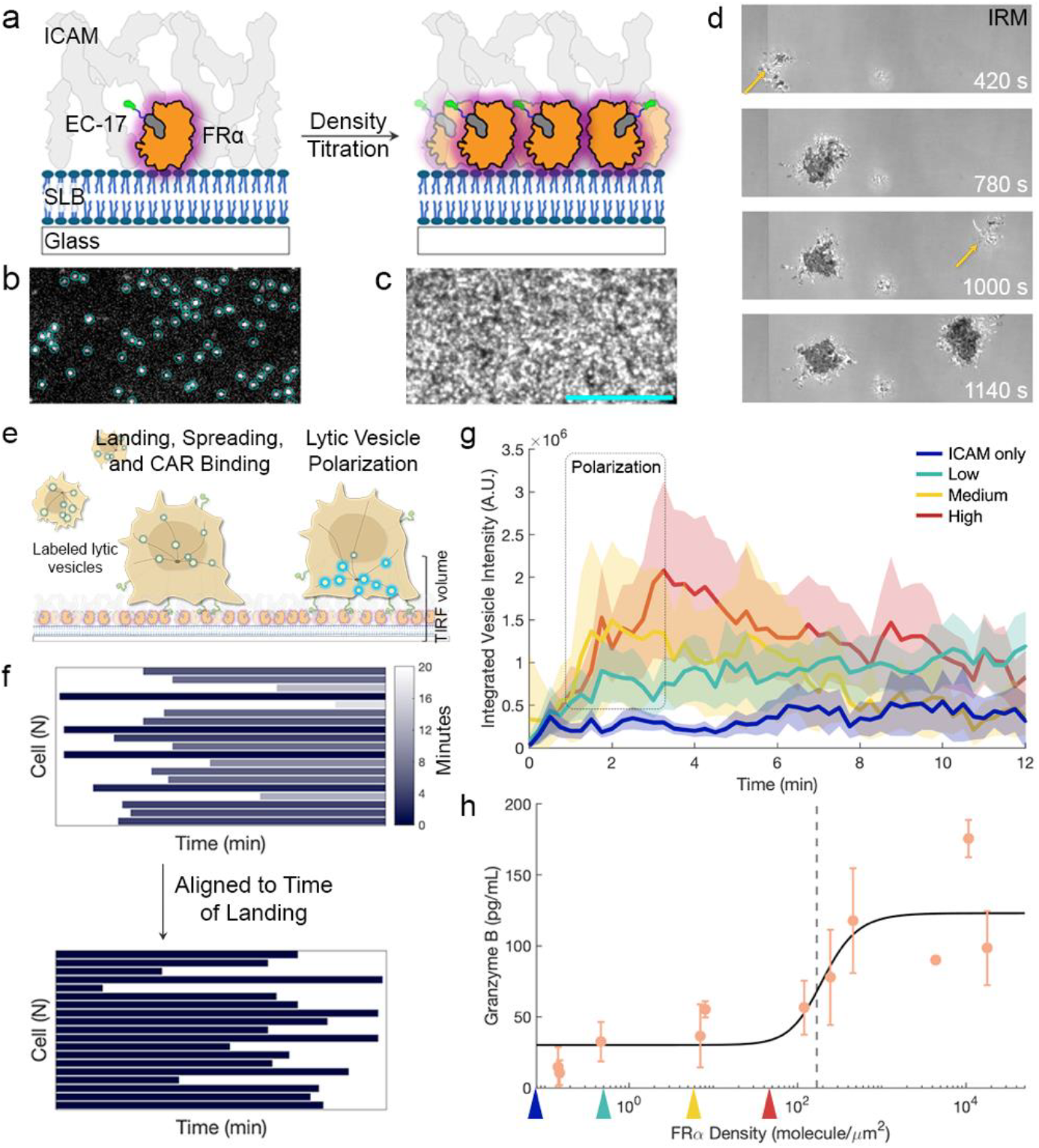
Population features of CAR T cell activation suggest a high agonist density threshold. **a.** *In vitro* reconstitution of CAR T cell:tumor cell junctions. FRα and ICAM ligands are displayed on the SLB surface at controllable densities. The density of FRα is varied over several orders of magnitude. **b-c**. Example images of fluorescently-tagged FRα displayed on SLBs at low and high densities, respectively. Shown are 0.10 and 22.70 molecules/μm^2^, respectively. **d**. Montage of IRM imaging of live CAR T cells landing on an SLB. Cells land, spread, and interact with the surface at different times. Initial contacts are indicated by yellow arrows. **e**. Raw tracks of CAR T cells interacting with an SLB surface based on the time of landing and the duration of interactions (top). Tracks are aligned to the time of landing, which is set as time = 0 (bottom). **f**. Schematic of the ensemble polarization assay showing that CAR T cells loaded with low pH marker for lytic granules interact with an SLB surface and, upon activation, undergo polarization and traffic the vesicles to the plasma membrane. **g**. For the aligned cell trajectories for each condition, the corresponding signal from the lytic vesicle channel was used to calculate an ensemble average trace for polarization. The shaded region around each trace is the 95% confidence interval. Data are from at least 150 cells in each condition from more than 3 biological replicates and multiple T cell donors. **h**. Supernatants were analyzed in a Granzyme B ELISA. Data are from at least 3 biological replicates at each agonist ligand density. Colored triangles correspond to the densities of FRα for the surfaces shown in (g).

We assayed population polarization using living CAR T cells loaded with the low pH sensitive dye, LysoBrite Blue, to mark the acidic cytotoxic granules laden with perforins and Granzyme B. Cells were allowed to engage SLBs and imaged for up to twenty minutes. Early landing and spreading interactions with the SLB were monitored using interference reflection microscopy (IRM; Fig. 2d) and polarization of the lytic vesicles was observed based on an increased and sustained fluorescent signal in the narrow TIRF volume near the plasma membrane (Fig. 2e). Across many fields of view, CAR T cells landed at different times and, thus, experienced different onsets of input accumulation (Fig. 2d; 2f; Fig. S2; Supp. Movie 2). The distribution of starting times for decision-making could obscure the fundamental activation mechanism. Thus, trajectories were aligned to landing, using the information from the IRM images (Fig. 2f)^40^. Across many tens of cells, rapid polarization occurred in response to high FRα densities of tens of molecules per square micron (Fig. 2g). At hundreds of molecules per square micron, the typical target expression levels of tumor targets, a similar rapid polarization phenotype was evident (Fig. S3). The fast rate of polarization matches prior reports for native cytotoxic T cells and CAR T cells^23,41,42^. As expected, there was very little polarization across the population of CAR T cells exposed to a surface displaying only ICAM adhesion ligands, emphasizing that signaling via the CAR was necessary for the response.

The combination of control over antigen ligand densities and ensuring the alignment of cell trajectories to time of landing enabled a more careful evaluation of CAR T cell behaviors at lower agonist ligand densities. Even at densities of less than one molecule per square micron, the population activation trace reflected a rapid increase in cytotoxic vesicles near the surface that was distinguishable from the non-stimulatory surface. The ensemble activation information suggests that some cells are robustly activating at low ligand densities despite an overall dampening of activation across the population. Activation responses were detectable as early as 2 minutes after landing. During the 20 minutes of the assay, cells that landed experienced long durations of contact and had ample time to experience enough inputs to permit polarization (Fig. S4).

Although mobilization of low pH vesicles in response to agonist ligand binding is a strong indicator of activation, we confirmed degranulation of cytolytic Granzyme B through an enzyme-linked immunosorbent assay (ELISA) using supernatants captured from the reconstitutions (Fig. 2h). As expected, high levels of Granzyme B release were observed at high ligand densities and the apparent threshold was set at hundreds of molecules per square micron^9,43^. However, it was also evident that some secretion occurred at low ligand densities of ones or fewer agonist molecules per square micron, similar to the polarization observed (Fig. 2g). At these low ligand densities, only small handfuls of binding events were possible due to rare encounters.

## CAR T cell activation at the single molecule level

CAR designs include cytoplasmic domains from the native T cell receptor and co-receptor signaling machineries and, thus, are expected to coopt the native signaling interactions and pathways to effect decision-making. However, most findings suggest that signal amplitudes are low and CARs ineffectively use key features of TCR signaling such as macromolecular assembly templated on the membrane tethered molecule Linker for Activation of T cells (LAT)^9,44^. To map signal amplitudes from single CAR binding events, we extended the reconstitution platform to detect each engagement between a CAR on a living T cell and FRα molecules loaded with EC-17. Single receptor:ligand complexes were unambiguously detected using a temporal filtering strategy that detects the slowed lateral mobility of ligands upon receptor engagement (Fig. 3a-b)^28,39^. The collection of binding events per cell provided spatial and temporal encoding of the inputs (Fig. 3c).

**Figure 3.**
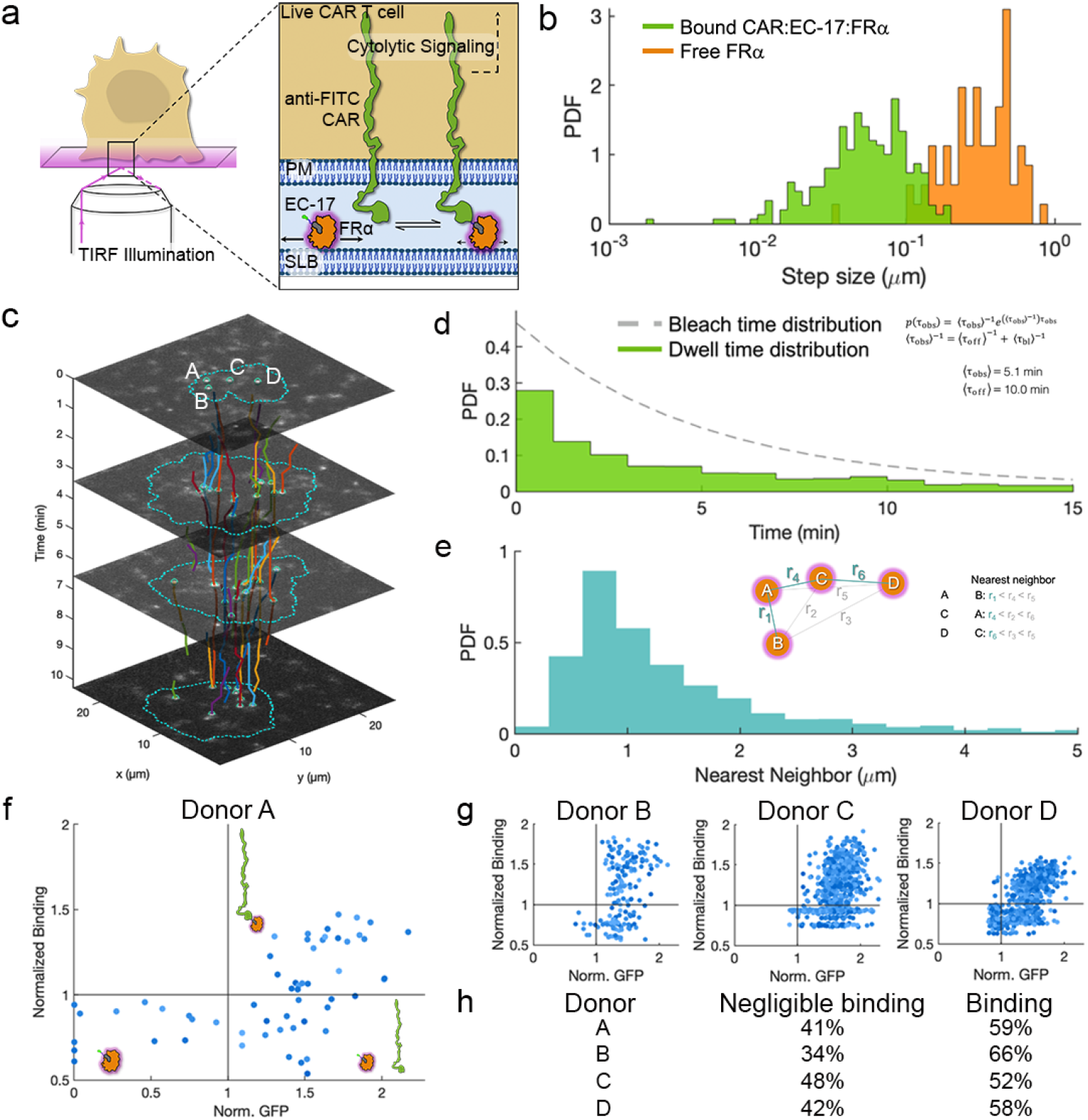
Single molecule binding of anti-FITC CARs to FRα generates long-lived, isolated binding events during information processing. **a.** Schematic of detection of single CAR engagement with antigenic ligand at the hybrid live cell-supported membrane interface. FRα on the SLB surface are singly labeled with fluorescent dye and loaded with EC-17 ligand and visualized using TIRF imaging. The ligands freely move with a fast diffusion coefficient until bound by a CAR on the T cell surface. **b**. Step size distributions for the mobilities of free FRα and bound ligands. The latter are shifted to smaller steps, consistent with a slower diffusion. **c**. Representative plot of CAR:EC-17:FRα trajectories collected by a single cell that was observed to land and collect binding events over a 10-minute span. Each track is colored and shown with x-y coordinates that reflect position and a z-coordinate for time. Labels A-D apply to panel (e). **d**. Dwell time distribution for CAR:EC-17:FRα binding events. The observed dwells and the bleaching curve are shown. **e**. Nearest neighbor distribution for binding events over the first 6-minutes after landing on a surface presenting <1 FRα/μm^2^. Inset schematic shows how nearest neighbors were determined using points A-D from panel (c) as an example. **f**. Binding probability for CAR T cells from Donor A. Data are plotted as intensity from CAR:EC-17:FRα binding interactions versus expression of GFP as a proxy for the CAR. Cells were exposed to the SLB surface for 20 minutes and endpoint data are plotted for accumulated binding. Schematics in each quadrant represent non-CAR-expressing cells (lower left), CAR expressing cells that experience negligible binding (lower right), and CAR expressing cells that binding (upper right). **g**. Binding probabilities for Donors B-D using the paradigm in panel (f). EC-17:FRα surface density for (f) and Donor B was tens of molecules per square micron and for Donors C-D was hundreds of molecules per square micron. **h**. Table of fraction of CAR-expressing, GFP-positive, cells that experienced binding for Donors A-D.

Biophysical features of CAR T cell signal accumulation can be extracted from the single molecule trajectories. The molecular binding dwell time was extracted from the durations of observed tracks. The nanomolar affinity of the anti-FITC scFv generated the long binding interactions that were expected estimated to be at least ten minutes (Fig. 3d). Since the typical timing of CAR T cell polarization was only a few minutes, all of the binding that contributed to activation outcomes were detectable throughout the relevant time and once bound, receptors typically remained bound throughout the signal integration time. We assessed the spatially proximities of CAR:EC-17:FRα complexes during the activation window of up to 6 minutes. At the high engagement densities that are predicted for CAR binding to TAAs, clustering is expected even at early times and some data argue that at CAR dimer is the minimal signaling unit^45^. At low EC-17:FRα densities where activation was detected (Fig. 2g), the distribution of distances between bound receptors was centered on almost one micron (Fig. 3e). Therefore, the signaling of isolated, ligated CARs must be effectively integrated to generate the polarization response without the need for other ligated receptors in the vicinity.

We noted that some cells collected negligible numbers of binding events despite forming large contact interfaces (Fig. S5). CAR T cells from four donors were exposed to SLBs presenting high densities of EC-17:FRα. After 20 minutes, the accumulated CAR binding was assessed. Using GFP signal as a proxy for CAR expression at the surface (Fig. S6), extent of binding did not purely scale with receptor expression (Fig. 3f-g). CAR T cells with negligible binding were observed across all donors and one donor population showed nearly 50% of CAR-expressing cells bound ligand negligibly (Fig. 3h). Thus, a subset of the CAR T cells that could respond to the available ligand are capable of both binding and activation.

To understand how single molecule binding at the intermembrane junction leads to CAR T cell activation, we developed a single molecule impulse-response (I-R) assay to map CAR:EC-17:FRα binding sequences to polarization outcomes at the single cell level. The I-R assay distills the complex signal integration process to the molecular transformation of individual impulses to a downstream, global response (Fig. 4a-b). This approach was previously applied to map TCR:pMHC biding to translocation of the transcription factor, nuclear factor of activated T cells (NFAT). The I-R assay demonstrated that the native TCR relies on spatial and temporal coordination of binding events to satisfy the long-engagement activation threshold^46^. Subsequent work identified molecular assembly based on the protein Linker for Activation of T cells (LAT) as the machinery for cooperativity^30^. Importantly, I-R assays detect the entire binding history and, by also monitoring the membrane interface, ensure that all data are recorded starting with landing and spreading on the SLB^16,30^.

**Figure 4.**
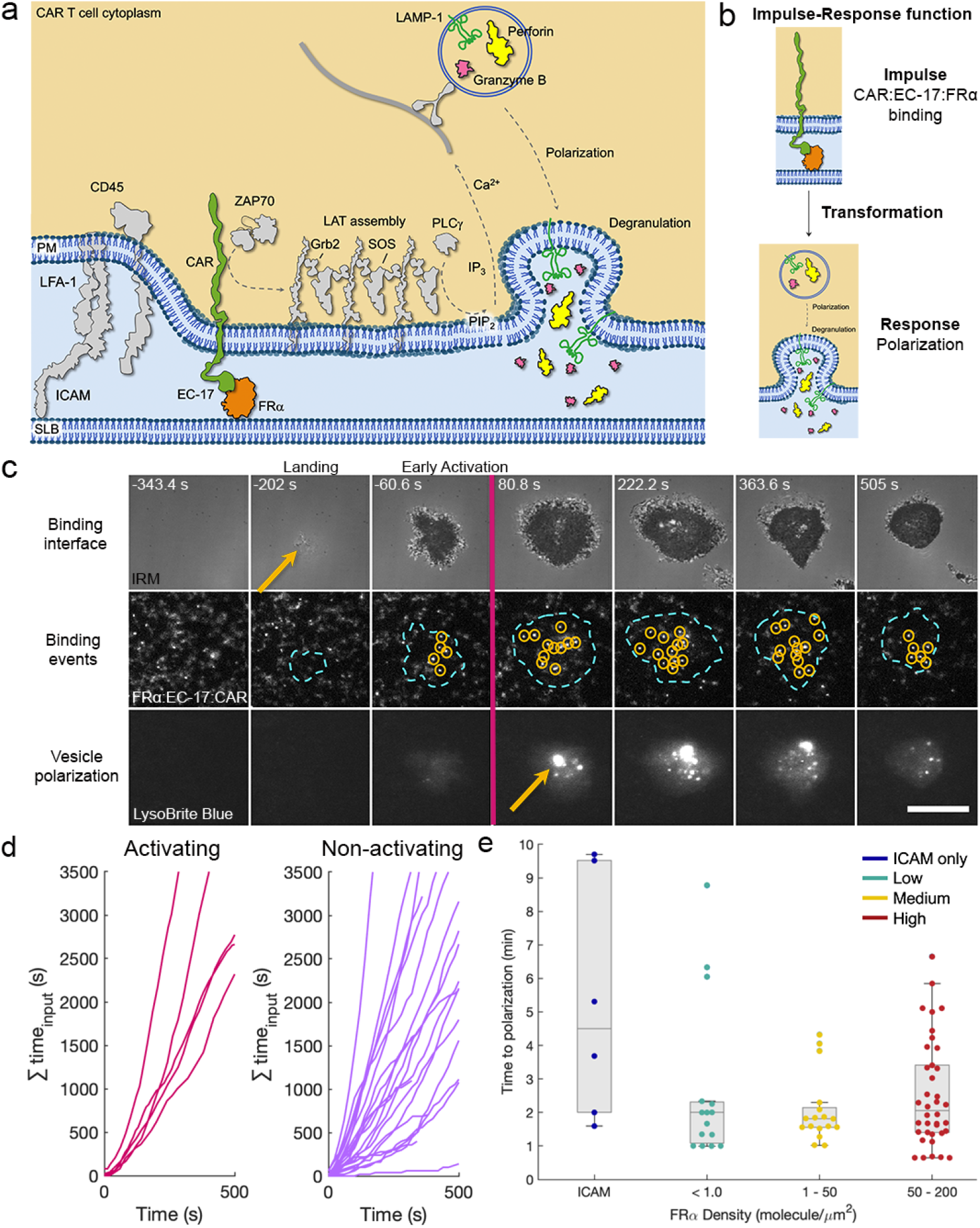
Molecular Impulse-Response (I-R) function shows that CAR triggering, at the single molecule level, sets the molecular activation threshold. **a.** Schematic of proposed intracellular signaling pathway that relays CAR:EC-17:FRα binding events to polarization and degranulation outcomes. The signaling pathway is based on the native cytotoxic T cell machinery. **b**. The signaling pathway is distilled into an impulse-response (I-R) function that monitors the individual binding events between CAR:EC-17:FRα (impulses) and maps the binding sequences to polarization of vesicles laden with cytotoxic molecules (response). **c**. Example I-R sequence at a FRα density of 0.82 μm^-2^. Montage of cell landing and spreading (IRM; top), impulse binding events (middle), and polarization response (bottom). The time of activation is set based on the detection of polarization. Scale bar = 10 microns. **d**. Accumulation of binding time for individual cells that either polarized or failed to activate. Each line represents the integrated binding time for a single cell. **e**. Single cell distributions for cells that polarized on ICAM only, and in low, medium, or high FRα density surfaces, respectively. Each point on the scatter represents a single cell. Data in (d-e) were collected from at least 4 independent experiments with CAR T cells from at least 3 different donors.

Polarization is the penultimate step in generating cytotoxic responses. The Granzyme B ELISAs already demonstrated that degranulation was occurring, but we wanted to verify this process in the I-R workflow. The process of lytic granule fusion occurs over a few hundred milliseconds^47,48^. A T cell will not exocytose its full payload and, instead, will detach from a target after delivering some cytotoxic granules and may subsequently generate serial toxicity. Rapid imaging of the fluorescently-loaded cytotoxic vesicles did, indeed, show degranulation events (Fig. S7; Supp. Movie 3). Given the rarity of these events, predicted to be around 3 degranulation events per cell, we focused on polarization in the I-R function^47,49^. Most I-R assays were conducted at low ligand densities of less than one molecule per micron square to ensure detection of the full binding history. The polarization probability was low when compared to higher agonist densities (Fig. S8) but some cells were observed to collect small handfuls of binding events and to rapidly activate (Fig. 4c; Supp. Movie 4).

The population of cells was divided based on collecting binding events and the binary activation readout of polarization. Accumulation of binding time did not distinguish activating and non-activating cells, arguing that those cells that polarize satisfy the activation threshold through a mechanism other than simply integrated interaction time. There was a slight distinction between the accumulated number of binding events (Fig. S9). However, some non-activating cells acquired as many binding events as activating cells suggesting that this parameter does not set the activation setpoint either (Fig. S10).

We measured the I-R function across FRα densities spanning three orders of magnitude. Some CAR T cells spuriously polarized on an ICAM-only surface but remained a low probability event. Remarkably, for cells that polarized on surfaces presenting FRα ligands, the mean time of polarization was centered on approximately two minutes and independent of ligand density (Fig. 4e). Therefore, activating cells were distinguishable through the rapid conversion of the first few binding events into the digital polarization response. These data demonstrate that mechanistic setpoint for CAR T cell activation is, in fact, very narrowly tuned and resembles that of the native TCR^14–16^.

## Crosstalk between the native TCR and CARs

The spacing between ligated CARs that we observed following landing argues for efficient propagation by single CAR:EC-17:FRα complexes using the native T cell machinery. The current engineering strategies in generating CAR T cell populations do not necessarily include alterations to the native TCR expressed on the T cell populations and could permit crosstalk between the receptors. This was recently highlighted by studies that showed TCR-CAR crosstalk that was tuned by the affinity of the TCR ligand^50^. In the cells used in the I-R assay, CAR expression did not alter TCR expression (Fig. S11). The presence of the TCR on CAR T cells interacting with surfaces that do not include peptide major histocompatibility complex (pMHC) ligands would be expected to remain inert. However, the presence of the additional 7 immunotyrosine activation motifs (ITAMs) on the endogenous TCR could contribute to the signal propagation by the CAR, which has only 3 ITAMs^51^. We note that the CAR used here undergoes ligation at a closer intermembrane distance than the native TCR which may correlate with exclusion of phosphatases as a mechanism to promote signaling^52^.

To interrogate a role for the TCR in modulating the CAR T cell impulse-response function, we employed a recently-introduced, transmembrane peptide inhibitor of the TCR (PITCR). The PITCR was developed based on the transmembrane sequence of the zeta chains of the TCR and is predicted to alter the orientation of these chains and diminish signaling by the TCR:pMHC complex (Fig. 5a;^53^). We integrated the PITCR into the ensemble polarization assay workflow by pretreating the CAR T cells with the peptide prior to exposure to an SLB presenting a high FRα density. A non-perturbative mutant, PITCR G41P, that showed no diminishment of TCR signaling, was also used^53^. The G41P and PITCR peptides led to trends that strongly resembled the WT CAR and ICAM-only polarization response curves, respectively, without altering the contact areas between the CAR T cells and the surface (Fig. 5b-c; Fig. S13). These are the first indications that the unligated TCR modulates CAR signaling amplitudes and argues that the single molecule sensitivity of the latter may be conferred by the endogenous TCR.

**Figure 5.**
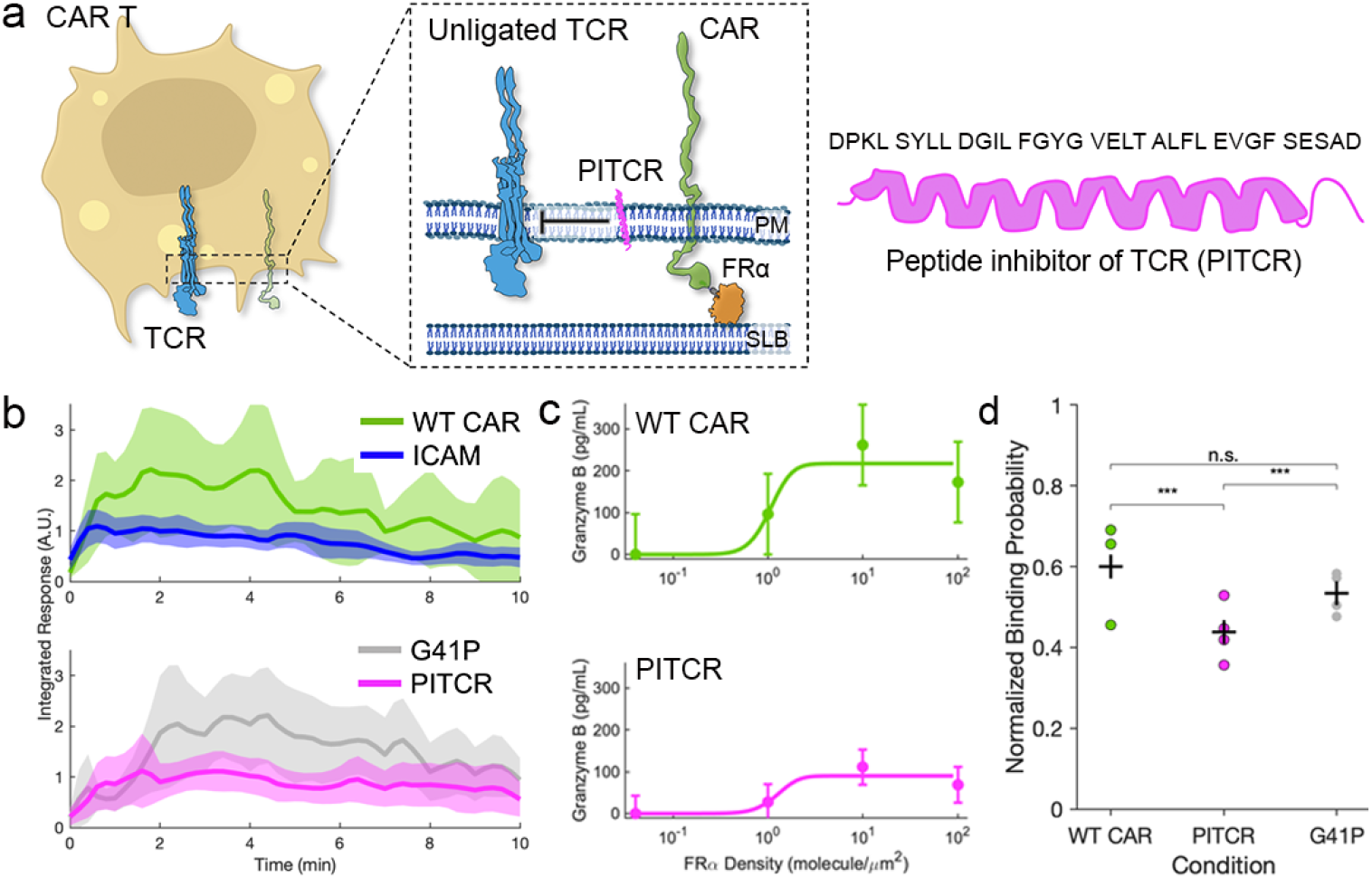
Disruption of the non-ligated TCR by a peptide inhibitor impacts CAR T cell signaling and activation. **a.** Schematic of CAR T cells, co-expressing native TCR, exposed to peptide inhibitor of TCR (PITCR). The PITCR will embed in the plasma membrane and was designed to block TCR signaling. **b**. Ensemble polarization of CAR T cells exposed to surfaces displaying high density EC-17:FRα (100s molecules per micron square), ICAM only (left), G41P, or PITCR (right), respectively. Traces are averages for cells aligned to landing. There are at least 50 cells in each condition from 3 biological replicates. Shaded region around each trace represents 95% confidence interval. **c**. Supernatants from reconstitutions were captured and run on a Granzyme B ELISA (WT CAR, top and treated with PITCR, bottom). Data are from at least 2 biological replicates at each agonist ligand density. **d-e**. Binding probability for CAR T cells from Donor D treated with PITCR or G41P transmembrane peptides, left and right, respectively. Data are plotted as intensity from CAR:EC-17:FRα binding interactions versus expression of GFP as a proxy for the CAR. Cells were exposed SLB surfaces with EC-17:FRα densities of hundreds of molecules per micron square for 20 minutes and endpoint data are plotted for accumulated binding. **f**. Summary of frequency of CAR-positive cells binding EC-17:FRα ligands. Each data point represents one biological replicate of at least tens of cells. Statistics are based on Chi squared test.

A mechanistic connection between the TCR and the CAR has been suggested based on observations that expression and signaling by the TCR can alter CAR expression and activities^50^. However, a role for the unligated TCR in modifying CAR T cell activities remains limited. Based on the findings using PITCR, we predicted that the endogenous TCR was modulating the signal propagation from the CAR. However, this could occur either at the level of successful CAR triggering or propagation following ligation. Given our finding that some CAR T cells experienced defects in ligand binding, we pursued a role for TCRs in modulating CAR binding. Introduction of PITCR significantly diminished CAR binding probability with more than 50% of cells binding ligand negligibly (Fig. 5d). These data suggest that organization of the plasma membrane, alone, modulates CAR binding but that the impact of the unligated TCR, specifically, has a marked effect. Regulation of the binding probability could be due conformational regulation of the CAR, organization withing in the membrane, or physicochemical regulation that controls the close intermembrane spacing necessary for receptor engagement.

## Discussion

Here, we show that CAR T cell activation hinges on a limited number of receptor:ligand binding events that are collected within the first few minutes of contact with the target and that this phenotype is restricted to a subset of the population. Thus, CAR T cells are capable of a rapid analog-to-digital conversion and satisfy the activation threshold criteria at agonist densities two orders of magnitude lower than the population ensemble and demonstrate an exquisite sensitivity in ligand detection and signal propagation. The ability of cells to encode stochastic information into binary responses has been demonstrated in gene regulation and is a hallmark of multiple stages of normal T cell activation^16^. Our impulse-response assay shows that mechanistically, CAR T cell polarization is the consequence of robust engagement of the endogenous signaling machinery and is promoted by the native TCR. The influence of the unligated TCR motivates new consideration of the activity of the endogenous receptor; coupled to other findings, there may be a spectrum of responses that are generated by the TCR based on the engagement of ligands and their affinities^50^. Normal T cell development and maintenance in circulation depends on signaling by the unligated TCR or in response to non-agonist-loaded MHCs. By comparison, tonic signaling from CARs may, paradoxically, drive improved antitumor signaling or promote T cell exhaustion, resulting in impaired function^54^. How the crosstalk between the unligated TCR and CAR regulates tonic signaling and the strength of antigenic responses will be an important direction for investigation.

CAR T cell therapies have been hampered by defects in infiltration and activation, and excessive damage to healthy tissues due to off-target toxicities. Deficits in activation are likely the result of stimulation not being restricted to tumorigenic cells given that the artificial signaling is not restricted to high ligand densities. Promising new technologies have focused on generating engineered receptors that signal more like the native TCR, including activating at low agonist ligand densities and engaging the endogenous signaling machinery^50,51^. These receptors have demonstrated potency against agonist ligands on tumors that are present at low densities and could prevent antigen escape. The dichotomy that antigen escape and normal tissues could be indistinguishable could be a significant impediment to the application of engineered cells in treating disease. An important recent finding in the context of chronic lymphocytic leukemia was that a limited number of CAR T cell clones could be isolated in patients with durable remission of more than a decade^1^. This is consistent with our findings that some cells are specialized in collecting binding events and predicts some of these cells generate sustained responses and persist in circulation. Additionally, CD4^+^ CAR T cells were shown to persist over many years, consistent with other findings that CAR T cell populations from each of the CD8^+^ and CD4^+^ lineages have differential potencies^1^. These populations were not distinguished in our study but could be characterized within the workflows we present. Collectively, our findings offer a new lens through which we should consider optimization of immunotherapies in T cells and other immune cells.

## Acknowledgments

We appreciate members of the Low-Nam lab for critical feedback and Tanvi Breinig for assistance with primary T cell culture. We acknowledge assistance from Ryan Schuck at UT Knoxville for providing the PITCR reagents. This work was partially supported by NIH grant R35GM140846 (to F.N.B).

## Materials and Methods

### Reagents

Folate Receptor α-10x His (Sino Biological) was conjugated with Alexa Fluor 647 NHS-ester (Fisher Scientific) using a 3:1 dye/protein molar ratio in 1 mM carbonate buffer (pH 10.05) for 20-30 minutes at 37 °C. The reaction was quenched with 20 molar excess Tris buffered saline. Free dye was separated using a 7 kDa molecular weight cut-off desalting column (0.5 mL, Zeba) and the protein conjugate was buffer exchanged to PBS (pH 7.4). The labeling ratio was measured using a NanoDrop (Fisher Scientific). The reaction was optimized for a final labeling ratio of 1:1. The labeled protein was aliquoted in single-use volumes, frozen in liquid nitrogen, and stored at -80 °C until used.

Loading of FRα with EC-17 was carried out 12-18 hours prior to imaging and performed fresh for each experiment. EC-17 at 125:1 molar excess (AdooQ Bioscience) was loaded onto FRα-AF647 at 37°C in PBS buffer (1x). When derivatizing the FRα-AF647-EC-17 complex onto SLBs, excess EC-17 was rinsed away before imaging the SLB.

Antibodies used were αCD3ε (clone OKT3, BioLegend 317346, PE/Dazzle conjugated), αCD8 (clone OKT8, LSBio LS-C764288-20, AF647 conjugated), αCD4 (clone OKT4, BioLegned 317422, AF647 conjugated), α FRα (clone 548908, R&D Systems 548908, AF488 conjugated), αICAM1 (clone MEM-111, NovusBiological NB500-318AF488, AF488 conjugated)

PITCR and G41P peptides were shared by the Barrera lab and are based on a prior publication^53^ Briefly, peptides were synthesized by Thermo Fisher Scientific (Waltham, MA, USA) and sequences were confirmed by mass spectrometry and reverse-phase high-performance liquid chromatography (HPLC). The PITCR sequence is DPKL SYLL DGIL FGYG VELT ALFL EVGF SESAD.

### T Cell Culture and Lytic Granule Labeling

CAR T cells were prepared by lentiviral transduction in the Low lab according to published protocols^26,55^. Cell populations consisted of approximately 50/50 mixtures of CD8/CD4+ primary human T cells. Periodically, CD8/CD4 ratios were monitored using αCD4 and αCD8 antibody flow cytometry and remained roughly 50/50 through the duration of culture. The 2^nd^ generation CAR construct was composed of anti-FITC CAR T cell domain (scFv from antibody clone 4M5.3, K_d_ = 270 fM), CD8α hinge and transmembrane domain, 4-1BB costimulatory domain, and CD3zeta domain^55^. CAR T cells were cultured in TexMACs GMP phenol-free medium (Miltenyi Biotec) supplemented with 100 IU/mL IL-2 (Miltenyi Biotec) and 2% human serum (Millipore Sigma). Cell passage occurred every 2-3 days to exchange media and maintain cells at a density of 1-2 million cells/mL. The day before imaging, CAR T cells were exchanged into IL-2 free complete TexMACs medium. Prior to imaging, CAR T cells were centrifuged (300xg, 4 min, 4°C) and the culture media was exchanged to PBS (1x) containing LysoBrite Blue (AAT Bioquest) at final 2x concentration. The cells were incubated at 37°C 5% CO_2_ for 30 minutes. Three washes were performed to remove excess label and exchange to live cell imaging buffer (LCB; 1 mM CaCl_2_, 2 mM MgCl_2_, 20 mM HEPES, 137 mM NaCl, 5 mM KCl, 0.7 mM Na_2_HPO_4_, 6 mM d-glucose, and 1% m/v BSA). Cells were stored in the incubator until imaged and used within 2 hours of LysoBrite labeling.

### Sample and Bilayer Assembly

Glass-supported lipid bilayer (SLB) membranes were prepared in AttoFluor imaging chambers and functionalized with proteins using standard protocols^56^. Small unilamellar vesicles (SUVs) were formed via extrusion (Avanti Polar Lipids) through a 0.1 µm pore polycarbonate filter (Cytiva). The lipid composition was 96 mol % 1,2-dioleoyl-sn-glycero-3-phosphocholine (DOPC) and 4 mol % 1,2 dioleoyl-sn-glycero-3-[(N-(5-amino-1-carboxypentyl) iminodiacetic acid) succinyl] (nickel salt) (Ni^2+^-NTA-DOGS) (Avanti Polar Lipids) in Milli-Q water (EMD Millipore). Round coverslips (#1.5 high tolerance, 25 mm diameter, Warner Instruments) were ultrasonicated (40 kHz) in 1:1 v/v isopropanol:H_2_O, rinsed then sonicated in 2% (v/v) Hellmanex III (Hellma Analytics) and etched for 5 minutes in piranha solution (3:1 v/v H_2_SO_4_:H_2_O_2_ Millipore Sigma). SUVs in 1x PBS were pipetted onto etched coverglass mounted in the imaging chambers and mixed via repeated pipetting, and bilayers were allowed to form via vesicle rupture for 10 minutes. Bilayers were blocked with 1% m/v bovine serum albumin (BSA; Sigma) for 20 minutes and rinsed with 1x PBS. To derivatize the surface, bilayers were incubated for 60 minutes with a range of solution concentrations (10-50 pM for single molecule studies, and up to 50 nM during stimulus titrations) of FRα-AF647-EC-17 and 2-5 nM human ICAM-1-10x His in 1x PBS. FRα-AF647-EC-17 densities ranged from <1-100s molecules/µm^2^. ICAM densities were 50-200 molecules/µm^2^. Just before imaging, bilayers were rinsed at least 3 times to exchange to live cell imaging buffer.

### Microscopy Setup

A Nikon Ti2 microscope with 4 laser lines (405, 488, 561, and 640 nm), motorized TIRF arm, epi-lamp arm, and live-cell incubation stage was used. At the start of each imaging day, the TIRF evanescent field was aligned by first centering the laser along the optical path using a back aperture camera and triangulation of the center point using 8 or more points sampled throughout the perimeter of the back aperture. Then the critical angle was defined by mapping TIRF angle to the intensity of sparse 100-nm Tetraspeck beads (ThermoFisher) on clean coverglass coated with 1 µg/mL poly-L lysine in 1x PBS. The peak detected intensity was selected for the critical angle for buffer solution (refractive index n = 1.33) and an additional fixed distance was added to account for cellular refractive index n = ∼1.38. That fixed distance was determined using SK-MEL2 cells expressing genome-edited clathrin-RFP and dynamin-GFP (gift from the Drubin Lab, UCBerkeley) and confirmed by the absence of scattered laser in T cell experiments.^57^ Most experiments were conducted using p-polarization of the TIRF field or a 50/50 mixture of p and s-polarization. Polarization was verified using a planar SLB labeled with DiD and mapped to back aperture camera.

### Live cell imaging

A live-cell incubation stage (Tokai Hit) was used to maintain an environment of 37°C and 5% CO_2_ for all CAR T cell measurements. For cell-cell killing assays, 50,000 MDA-MB-231 cells were plated onto coverslips and allowed to spread for at least 10 hours in an incubator. Target tumor cells were well-separated to enable detection of each contact with CAR T cells. Cells were transferred aseptically to an imaging chamber and equilibrated in LCB. The chamber was mounted on the microscope. Time zero for a cell:cell killing experiment was the time at which EC-17, at a final concentration of 100 nM, and CAR T cells, at a 4:1 effector:target (E:T) ratio were simultaneously added to the chamber, in suspension. Imaging was performed using DIA and IRM channels for 8 hours, with a time resolution of 3 minutes. We performed 3 biological replicates to select a total of 100 target cells for which contacts and killing profiles were curated.

The SLB-containing imaging chamber was pre-warmed for 5-10 minutes before images of FRα-AF647-EC-17 density were taken in a peripheral region of the SLB. CAR T cells were pre-warmed to 35°C to avoid focal drift upon addition to the imaging chamber. For each chamber, 0.5 million cells were added. Image acquisitions for impulse-response (I-R) data came in 2 general forms. 1. For ensemble polarization, tiled 7×3 (columns x row) fields of view (FOVs) imaging used three sequential channels (Interference Reflection Microscopy; IRM for the contact interface, TIRF for CAR:EC-17:FRα-AF647 binding, TIRF for LysoBrite granule polarization) at a 20-30 second interval for 20-30 minutes. A complete tile was acquired in each channel before acquiring the next channel (imaging hierarchy of time, channel, position) or 2. Single FOV imaging in the same three channels, with a 10-20 second interval and a duration of 8-15 minutes. The latter I-R sequence was occasionally acquired at 2-4 x-y, non-overlapping positions instead of a single FOV, using an imaging hierarchy to match the individual FOV (time, position, channel). IRM enabled detection of the initial CAR T cell contact with the cell as a small, dark feature with an area of at least 5 µm^2^. The TIRF CAR:EC-17:FRα-AF647 binding channel discriminated slowly diffusing, single-molecule binding events by using a low power (55 µW) and long exposure time (500 ms). TIRF LysoBrite granule polarization channel was used to visualize lytic granules were transported to the intermembrane interface, within the TIRF evanescent field of approximately 150 nm in depth.

After each I-R assay, images of the GFP marker for CAR expression and all other channels were acquired for FOVs imaged. The mobility of some CAR T cells required imaging in a larger tiled area than the original I-R assay. Occasionally, some additional endpoint images were acquired in a tiled fashion (3×3 or 5×5 FOVs) for cells not in the area of the I-R assay acquisition to sample additional cells. These data were analyzed independently from time-of-landing-aligned data. For some I-R assays, supernatants were collected to measure soluble molecules released into solution (expanded in the ELISA section).

### Flow cytometry

A BD Accuri C6 plus in the standard 3-blue (488 laser with 530/30, 585/40, and 670 LP filters) and 1-red configuration (640 nm laser with 675/25 filter) was used for flow cytometry profiling of CAR T cells. Fluorescence calibration beads for GFP signals (Takara) and AF647 (Bangs Laboratories) were used to convert arbitrary fluorescence counts into units of number of molecules per cell (corrections for individual antibodies were made to account for multiplicity of labeling).

### ELISA

To assay the ensemble levels of degranulation, CAR T cells were exposed to various densities of FRα on SLBs containing FRα and ICAM-1 for 30 minutes at 37°C. At 30 minutes, 800 μL of supernatants above the SLB were collected and spun down at 1000xg for 10 minutes to remove any cells or debris. The supernatant aliquoted and stored at -20°C. Supernatants were thawed and an ELISA kit for granzyme B (Invitrogen) was used to measure granzyme B levels.

### Estimating FOLR ligand Density

The number of FRα molecules on the SLB surface was estimated by relating the integrated fluorescence over a fixed area to the integrated intensity of well-isolated single molecules.

### Image Analysis

An analysis pipeline, implemented mainly in MATLAB (MathWorks, Natick,MA), was developed (available on GitHub) to process raw microscopy data. Raw data in the .nd2 format from Nikon Elements were loaded in MATLAB to extract metadata and timestamps. IRM or Diascopic images were corrected using a median filter^58.^ Fluorescence images were converted to EMCCD gain-corrected intensity values. TIRF images were flatfield corrected using shade images normalized collected on SLBs labeled with a carbocyanine dye (DiO, DiI, or DiD; ThermoFisher) and normalized using BaSiC^59^. Individual CAR T cells were isolated using manual selection of regions of interest (ROI) that delimited the overall area contacted during the acquisition. Cell segmentation was achieved using the IRM channel by using the prior that most bright pixels in the image represent background (bkgd), that bkgd is represented by a Gaussian distribution, and cell contact intensity thresholds are 3-5 standard deviations lower than the bkgd. Time of landing was captured from the IRM data to use in alignment of trajectories to first contact (as shown in Fig. 2). Manual curation of masks ensured that cells were fully separated. Semi-automated linking of masked images for a single cell in time was done using a Kalman filter and manually correcting any discontinuities.

Preprocessed data were saved and .TIFF and imported to ImageJ for tracking using the plugin TrackMate^60^. Single molecule puncta of bound FRα were localized using a Laplacian of Gaussians detector and quality threshold above the background, identified visually. Tracks were elongated using the Linear Assignment Problem Tracker with a 1 µm linking distance, and gap closing of 1 µm and max 5 frame gap. Tracks of only 1 frame in length were excluded from the dwell time distribution and bleaching rate calculations. Track coordinates were exported to MATLAB for further analyses.

LysoBrite-loaded, cytotoxic-vesicle polarization was determined by measuring the fluorescence intensity in the TIRF imaging volume. For each cell, the cellular background was fit using a Gaussian model. Polarization was defined as signal at least two sigmas above the mean background intensity.

