## Supplementary Figures for "CAR T cell cytotoxic responses are rapidly generated and sensitive to the unligated TCR"

**Supplemental Figures**

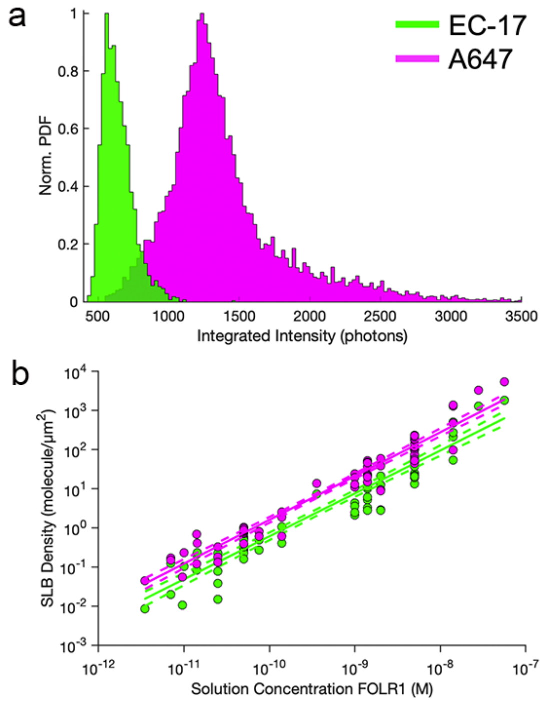

**Figure S1. Determination of surface FRα densities on supported lipid bilayers using single molecule intensities**. **a**. Single molecule intensity distributions for >2,500 single molecule localizations for each of FRα-AlexaFluor647 (A647; magenta) and EC-17 (FITC-folate; green) signals. **b**. Surface density of FRα measured by A647 or FITC fluorescence. Each point represents the average density of at least 10 fields of view (FOV) per SLB. Each FOV is cropped to the central 256x256 region of the image. Densities are calculated from individual localizations of well-resolved spots at low densities or scaled, based on average overall intensity divided by single molecule brightness, at higher densities. Same color scheme as in (a).

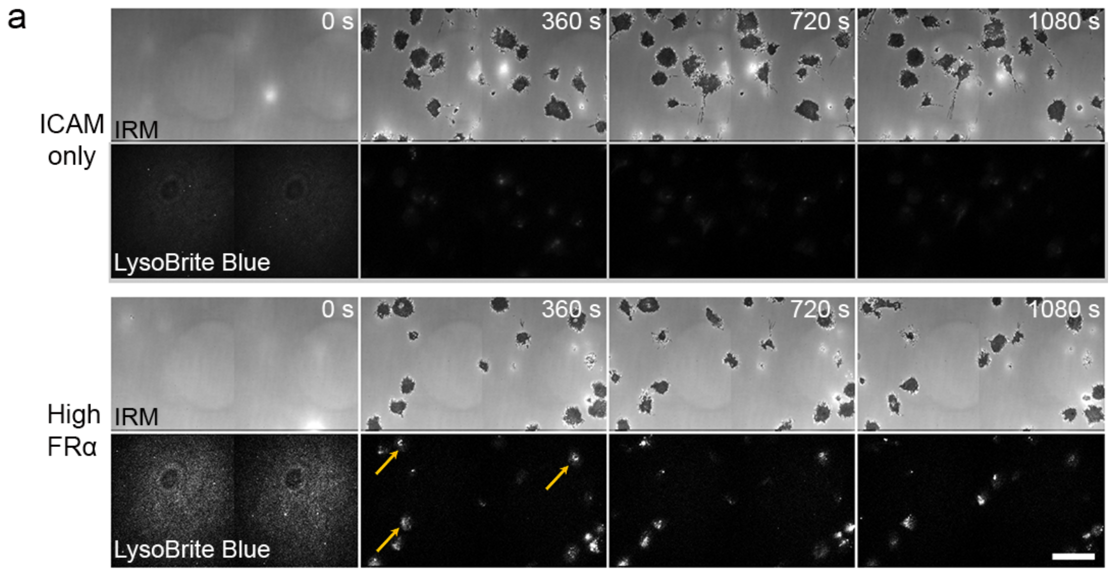

**Figure S2. Ensemble landing and polarization on different surfaces**. **a.** Example montages of CAR T cell landing, spreading in IRM, and polarization, indicated by recruitment of LysoBrite Blue to the interface. Surfaces are ICAM only (top) and high density FRα (116 FRα molecules/μm$^2$; bottom). Some polarized cells are indicated by yellow arrows. Scale bar = 20 microns.

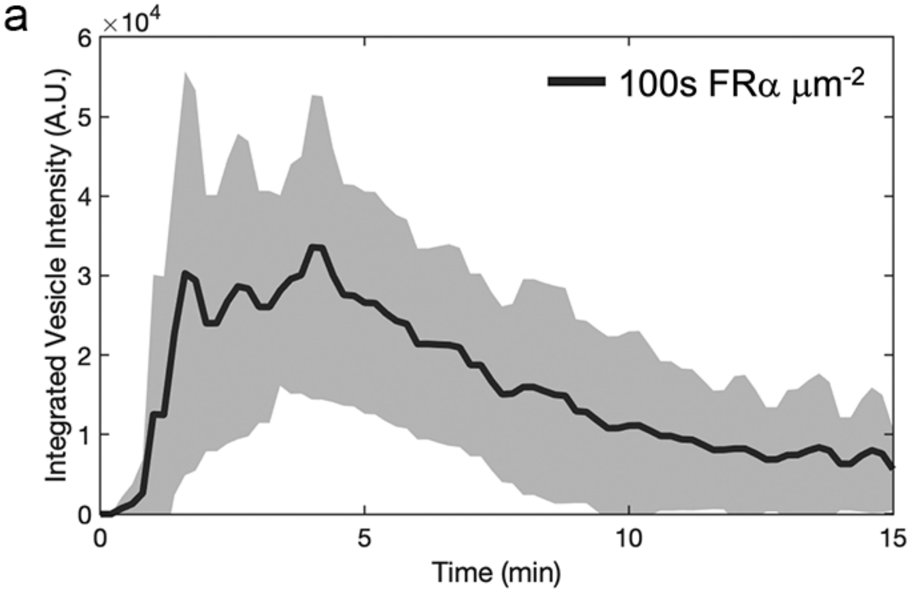

**Figure S3. Ensemble polarization of CAR T cells at FRα density of >400 molecules per micron square**. **a**. Using cell trajectories that are aligned to time of landing, the corresponding signal from the lytic vesicle channel was used to calculate an ensemble average trace for polarization. The shaded region is the 95% confidence interval. Data are from at least 150 cells in each condition from 2 biological replicates.

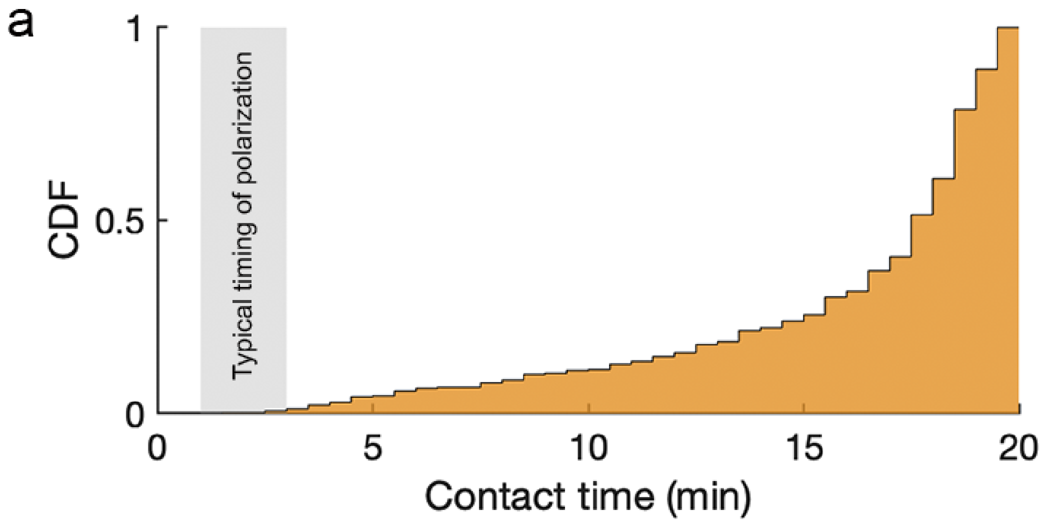

**Figure S4 – Cells that land on SLBs presenting ICAM and FRα maintain extended contact and have sufficient time to make activation decision. a.** Distribution of contact times during a 20-minute acquisition. Cell contact times are recorded from onset of landing and spreading. Data are aggregated across all surfaces and FRα densities and represent >1000 cells across tens of experiments and multiple donors. Shaded gray region shows typical time during which polarization is observed

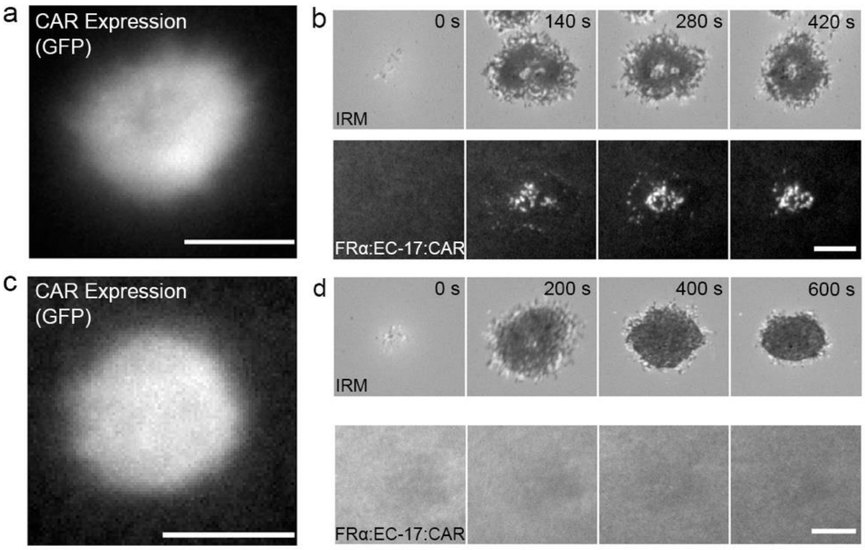

**Figure S5 – Some CAR T cells experience negligible agonist binding despite CAR expression and available FRα ligands. a.** GFP expression in CAR T cell corresponding to montage in (b). **b**. Landing and spreading in IRM (top) and binding events (bottom) at high FRα (>100s FRα molecules/μm$^2$). **c**. GFP expression in CAR T cell corresponding to montage in (d). **d**. Landing and spreading in IRM (top) without detectable binding (bottom) at the same high ligand density. Scale bar = 5 microns.

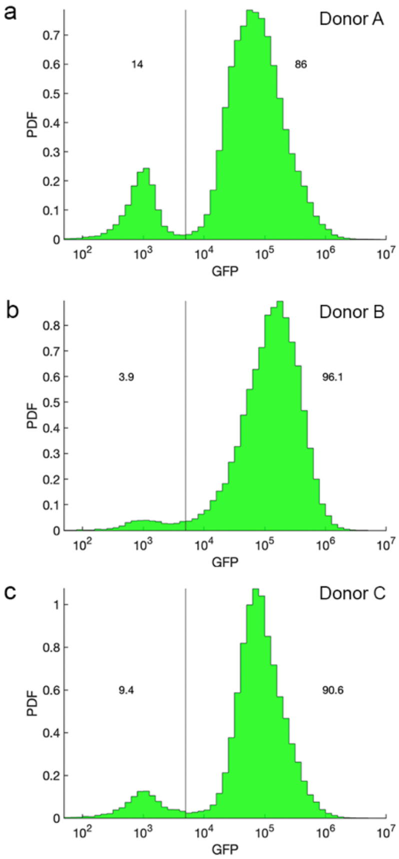

**Figure S6. Distribution of CAR expression levels**. **a-c**. GFP expression levels across 3 human donor (Donor A-C) T cell populations, measured by flow cytometry. At least 25,000 CAR T cells analyzed per donor. Vertical line shows gating for CAR positive cells and fraction expressing GFP are shown near the peak.

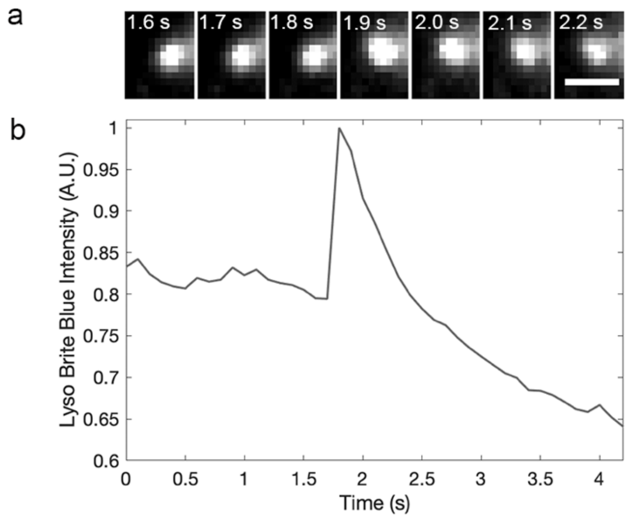

**Figure S7 – Direct observation of degranulation by CAR T cells. a.** Montage of Lyso Brite Blue signal at a spot undergoing degranulation. **b**. Intensity trace for the event shown in (a). The puff degranulation event occurs within hundreds of milliseconds. Scale bar = 1 micron.

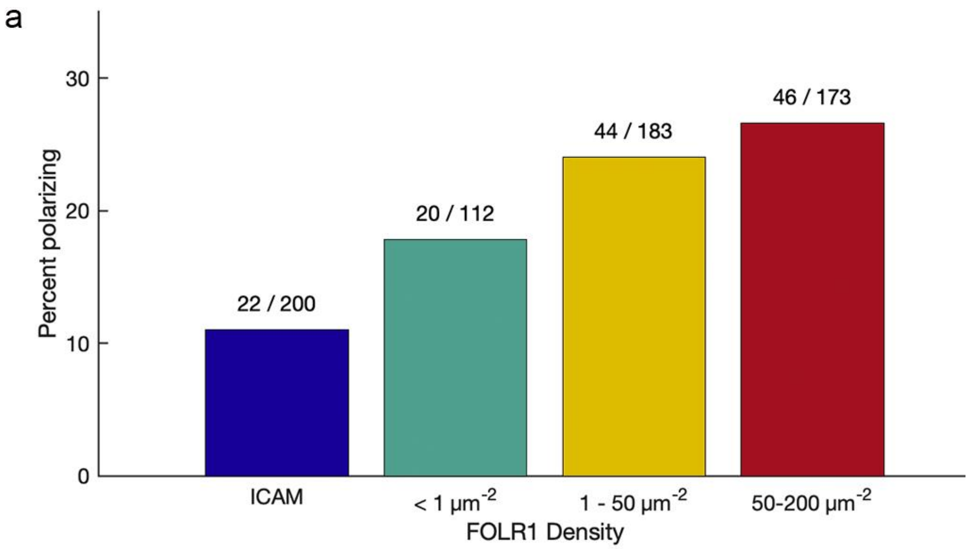

**Figure S8 – CAR T cells polarization probabilities scale with kinds of inputs and FRα density. a.** On each surface, cells observed to land were assessed for polarization. Cells with prominent uropod structures contacting the SLB were ignored due to the high probability of granule signal near the surface by virtue of the uropod structure alone.

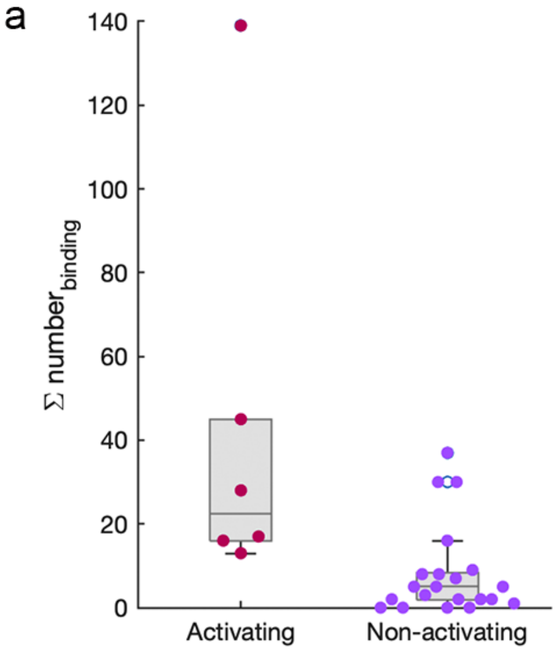

**Figure S9 – Integrated numbers of binding events between activating and non-activating cells at low, single molecule FRα density. a.** Single cell spreads of total number of binding events accumulated by cells on FRα at a density of 0.82 $\mu m^{-2}$. For non-activating cells, total binding is based on impulse accumulation up to the average time of activation for activating cells (~2.5 minutes).

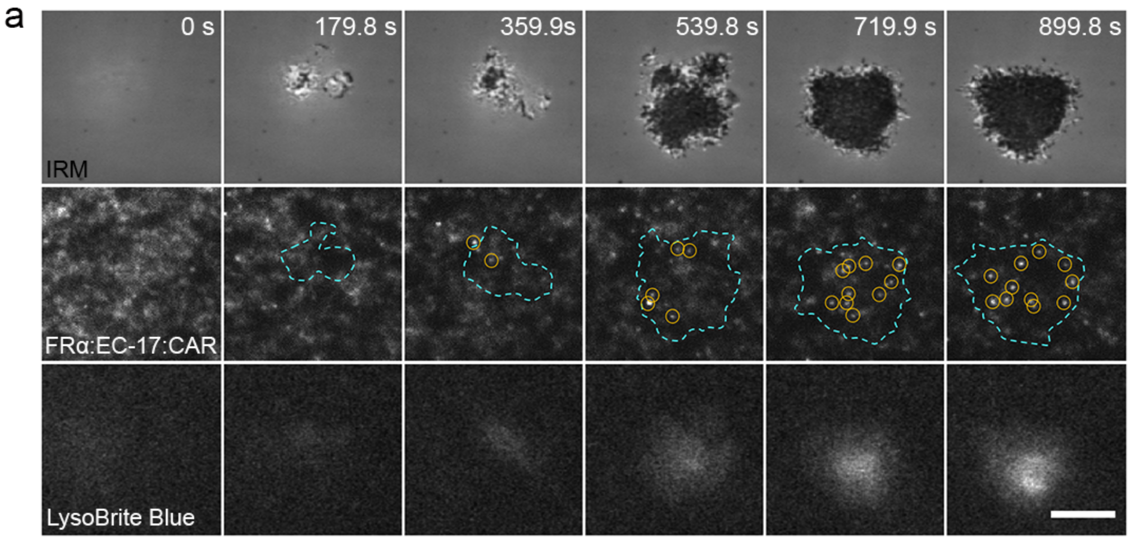

**Figure S10 - Example Impulse-Response montage for a non-activating CAR T cell at low FRα density (0.80 molecule/μm²). a**. Montage of cell landing and spreading (IRM; top), collecting impulse binding events (middle), but lacking a polarization response (bottom). Scale bar = 5 microns.

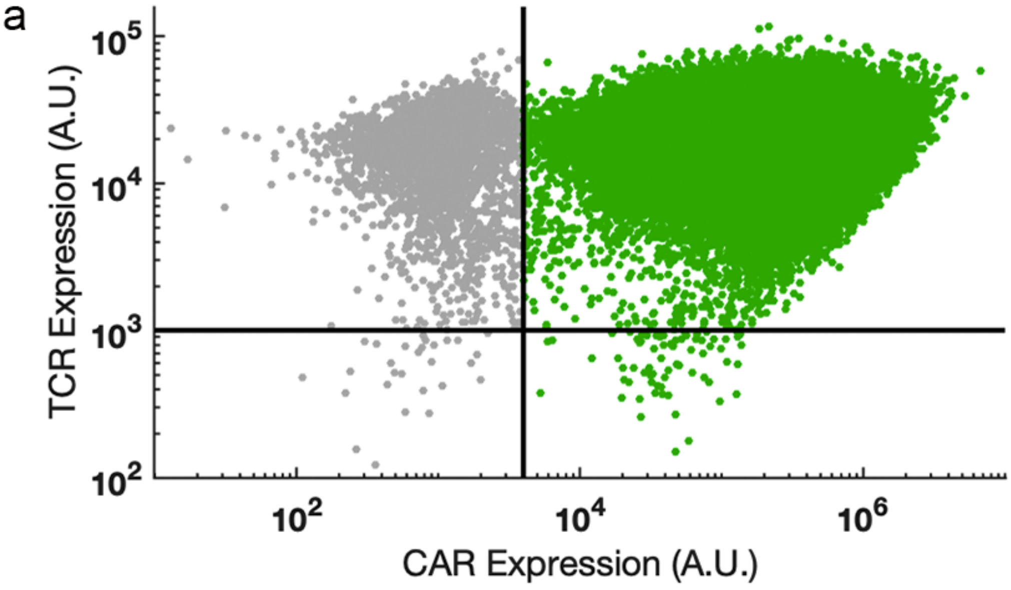

**Figure S11 – TCR expression levels are not impacted by CAR expression**. **a**. Anti-CD3 flow cytometry to detect TCR expression. TCR levels are plotted against GFP fluorescence as a proxy for CAR expression levels. Cells colored gray were gated as non-CAR expressing and those in green express the CAR.

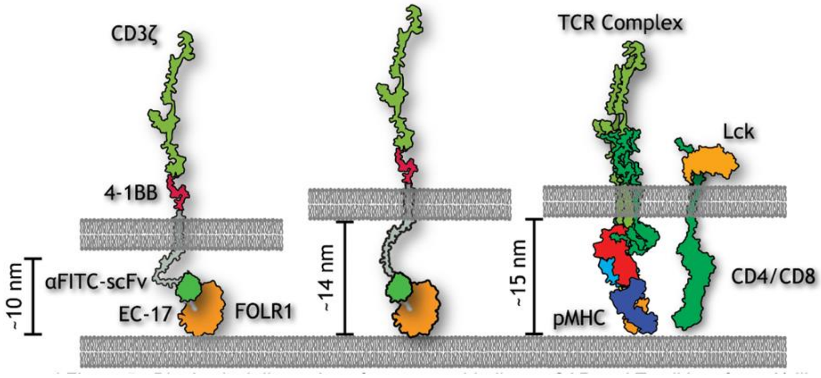

| Protein (domain) | PDB source |
|---|---|
| FOLR1 - folate bound | 4LRH |
| αFITC (scFv) | 1X9Q |
| CD8α (TM domain) | AF-P01732-F1-model-v4 |
| 4-1BB/CD137 (cytosolic domain) | AF-Q07011-F1-v4 |
| TCR-pMHC complex | 7PHR |
| CD3zeta (cytosolic domain) | AF-P20963-F1-model-v4 |
| CD3epsilon (cytosolic domain) | AF-P22646-F1-model-v4 |
| CD4 | AF-P01730-F1-model-v4 |
| Lck | AF-P06239-F1-model-v4 |

**Figure S12 - Biophysical dimensions for receptor binding at CAR and T cell interface**. A) Schematic of PDB crystal structure 2D projections for T cell receptor and CAR, their ligands, and some associated molecules at the membrane. Estimates of intermembrane distances reflect the flexibility of the extracellular hinge of the CAR.

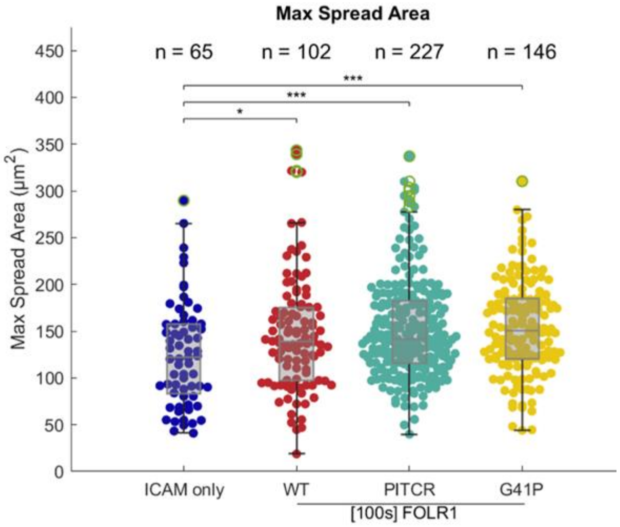

**Figure S13 - Maximum spread area for CAR T cells is not impacted by treatment with transmembrane peptides**. **a**. Single cell spreads for maximum contact area, measured in IRM, for cells on ICAM only or ICAM + high density FR$\alpha$ surfaces. Cells treated with the PITCR or G41P peptides spread as well as the WT cells. Wilcoxon rank sum test. '*' is p<0.5 '***' is p<0.01.

**Supplemental Movies**

**Supplemental Movie 1 – CAR T cell killing of target cell with minimal contacts**. Target MDA-MB-231 cell is spread on a glass substrate. CAR expressing cell (co-expressing GFP; shown in green) initiates contact and rapidly kills the target.

**Supplemental Movie 2 – CAR T cells land and spread on reconstituted surfaces**. IRM imaging of CAR T cell landing and spreading on SLBs presenting ICAM. Many fields of view are sampled using a tiling strategy. Scale bar = 50 microns.

**Supplemental Movie 3 – CAR T cell degranulation**. Example of degranulation event detected by rapid imaging of LysoTracker in TIRF (left). Magenta box indicates where the degranulation event occurs and provides the area over which the intensity trace (right) is recorded. Scrolling bar on plot provides the corresponding point in the raw data. Cell outline in cyan. Scale bar = 3 microns.

**Supplemental Movie 4 – CAR T cell impulse-response function at low FR$\alpha$ density**. Timelapse of a single CAR T cell interacting with an SLB with <1 FR$\alpha$/$\mu m^2$. Movie corresponds to montage in (4c). Landing and spreading are detected in IRM (left), binding event impulses are localized (middle) and the polarization response (right) is based on detection of loaded granules recruited to the surface. Scale bar = 5 microns.
